# Beyond Cell Cycle Control: *CDKN2A* Loss Orchestrates NAD^+^ Metabolic Plasticity and NAMPT Inhibitor Sensitivity in Glioblastoma

**DOI:** 10.64898/2025.12.02.691903

**Authors:** Swati Dubey, Guanqiao Yu, Ryana Aboul-Hosn, Christopher Tse, David A Nathanson, Albert Lai, Keith Vossel, Fausto J Rodriguez

## Abstract

While *CDKN2A* loss is classically associated with cell cycle deregulation through the p16-Cdk4-Rb axis, our findings suggest an additional layer of metabolic vulnerability arising from altered NAD homeostasis in *CDKN2A*-deleted glioblastoma, revealing a previously unrecognized metabolic-genetic interface for rationally revisiting NAD^+^ targeting strategies, moving beyond the broad inhibition approaches.

## Introduction

Despite extensive therapeutic advances, glioblastoma (GBM) remains one of the deadliest human malignancies, with median survival rarely exceeding 18 months. One of the defining hallmarks of GBM is its extraordinary metabolic plasticity enabling adaptation to diverse nutrient and redox environments. Distinct molecular subtypes of GBM display divergent metabolic preferences,^1^ however, the molecular circuits enabling these metabolic adaptations remain poorly defined. Among metabolic pathways supporting these adaptive programs, lies nicotinamide adenine dinucleotide (NAD^+^), which fuels multiple metabolic and non-metabolic pathways, including glycolysis, oxidative phosphorylation, fatty acid oxidation, and the activity of PARPs and sirtuins.^2,3^ Early enthusiasm for targeting NAD^+^ metabolism in cancer centered on inhibition of the rate-limiting NAD^+^ salvage enzyme nicotinamide phosphoribosyltransferase (NAMPT), which is overexpressed in GBM also and correlates with an adverse clinical outcome.^2^ Selective NAMPT inhibitors such as FK866 and KPT-9274 effectively depleted intracellular NAD^+^ and induced apoptosis across multiple tumor types, including GBM.^3^ However, dose-limiting toxicity in clinical trials dampened momentum and led to the assumption that therapeutically targeting NAD^+^ may not be feasible. Recent advances have revitalized this area. It is now clear that NAD^+^ metabolism is compartmentalized and context-dependent, with distinct regulatory mechanisms in the cytosol, nucleus, and mitochondria, and that cancer cells may rely on specific, rather than global, NAD^+^ dependencies. Recent reports, including the mitochondrial arm of NAD^+^ metabolism with the discovery of SLC25A51 as the principal mitochondrial NAD^+^ transporter, have begun to uncover new layers of complexity in NAD^+^ homeostasis and metabolic plasticity.^4,5^ In addition, a very recent study identifying gliocidine as a modulator of NAD^+^ dependent metabolism in GBM underscore a growing recognition that the NAD^+^ network is far more dynamic and context-dependent than previously appreciated.^6^ This resurgence underscores that instead of abandoning the NAD^+^ targeting strategies due to NAMPT inhibitor toxicity, selective and context-specific vulnerabilities rather than pan-cancer NAD^+^ depletion may hold the key to successful translation of NAD^+^ based therapies. Here, we revisited NAD metabolism as a therapeutic frontier in GBM, integrating transcriptomic analyses (TCGA-GBM dataset) with NAD^+^ perturbation studies using NAMPT inhibitors in GBM cell lines and identified a previously unrecognized link between NAD^+^ metabolic dependency and *CDKN2A* deletion, one of the most frequent genomic aberrations in GBM.

## Materials and Methods

### Cell lines and culture conditions

U251 (adult glioblastoma) and SF188 (pediatric high-grade glioma) were cultured in DMEM/F-12, GlutaMAX™ supplement (Gibco™ # 10565018) added with 10% FBS. JHH-NF1-GBM1 (NF1-associated glioblastoma line) and all the GS lines (Patient Derived GBM Cell Lines) were maintained in DMEM/F12, HEPES (ThermoFisher # 11330057) containing 2mM GlutaMax (Gibco™ #35050061), 1X B27 supplement (Gibco™ #12587010), 20 ng/mL EGF (PeproTech, Cranbury, NJ, USA), 20 ng/mL FGF-b (PeproTech), 5 µg/mL Heparin (Millipore SIGMA, Burlington, MA, USA) and 1X Pen-Strep (ThermoFisher # 15140122). TM31 (NF1-associated astrocytoma) was grown in MEM (Gibco #11095080) with 10% FBS, and normal human astrocytes (NHA) were cultured in DMEM (Gibco #11995065) with 2 mM GlutaMAX and 1X Pen-Strep supplemented with 10% FBS. All cultures were maintained at 37°C, 5% CO_2_. Cell identity was confirmed by STR profiling, and mycoplasma testing was performed routinely.

### Cell viability and dose-response curve

PrestoBlue assay was used to assess cell viability. Optimal seeding densities were 1,000 cells/well for U251, SF188, GS lines, and NHA, and 10,000 cells/well for JHH-NF1-GBM1 in 96-well plates. Cells were plated in 100 µL of culture medium and kept in incubator for 12-16 hours prior to drug treatment. Serial dilutions of candidate compounds GNE-617 and FK866 were added (final volume 200 µL/well) and incubated for five days. Following incubation, 20 µL of PrestoBlue reagent was added, and fluorescence (excitation/emission: 560/590 nm) was measured after 2 hours using Molecular Devices SpectraMax M5 microplate reader. Viability was normalized to DMSO controls, and dose-response curves were generated using nonlinear regression in GraphPad Prism.

### In silico analysis

All analyses were conducted in R (v. 4.3.2). TCGA-GBM gene-level CNV and RNA-seq data were retrieved using TCGAbiolinks R package (PMID: 26704973). CNV data generated through the ABSOLUTE LiftOver workflow were used to define CDKN2A alteration status. Samples with a maximum copy number of 0 were classified as CDKN2A deep-deletion group (DD), whereas those with copy number ≥ 2 were classified as non-deletion (ND). We then matched RNA-seq count data with the grouping method and ran DESeq2 (v. 1.4.6.0) (PMID: 25516281). Gene-set enrichment analyses (PMID: 16199517; 26771021) of DD vs. ND gene sets were performed to identify pathways affected by CDKN2A loss.

### Code availability

Scripts used to produce figures presented in this manuscript can be downloaded from GitHub at https://github.com/EdgeYu97/CDKN2A-GBM-Analysis

## Results and Discussion

We evaluated the sensitivity of a panel of GBM cell lines to two distinct NAMPT inhibitors, FK-866 and GNE-617. Both compounds markedly reduced proliferation across all GBM lines, whereas normal human astrocytes (NHA) maintained proliferation, indicating tumor selective vulnerability. Notably, most GBM cells harboring *CDKN2A* deletion showed heightened sensitivity to NAMPT inhibition compared to *CDKN2A*-intact counterparts (Fig 1a), which prompted us to investigate the link between *CDKN2A* status and NAD^+^ metabolism. To this, we analyzed TCGA-GBM transcriptomic datasets stratified by *CDKN2A* status. Tumors with *CDKN2A*-deep deletion showed significant enrichment of oxidative phosphorylation (OXPHOS) (Hallmark OXPHOS, NES = 2.57, padj <0.01) (Fig 1b), consistent with reports that RB pathway disruption drives oxidative metabolism and increases NAD^+^ demand.^7^ Then, we looked into the NAD^+^ salvage pathway. Although the overall NAD^+^ salvage pathway was not significantly enriched (REACTOME_NICOTINAMIDE_SALVAGE, NES = 1.7, padj = 0.1), individual genes showed distinct regulation: *NAMPT* was significantly upregulated (p<0.05), whereas *NMNAT2*, which converts NMN to NAD^+^, was downregulated (p<0.001) in *CDKN2A*-deleted tumors (Fig 1c, f). Given the elevated OXPHOS signature, we anticipated increased mitochondrial NAD^+^ import in *CDKN2A*-deleted tumors. Surprisingly, this oxidative phenotype coincided with significant downregulation of *SLC25A51* (p<0.001), the principal mitochondrial NAD^+^ transporter (Fig 1c). This downregulation of *SLC25A51* presents a paradoxical scenario and made us question, how do *CDKN2A*-null tumors sustain high oxidative metabolism under constrained NAD^+^ import capacity. One possibility is that the NAD and redox network in *CDKN2A*-deleted tumors might have rewired to sustain high OXPHOS and redox homeostasis. Interestingly, we found that this downregulation of *SLC25A51* was accompanied by upregulation of other mitochondrial transporters which play essential roles in shuttling metabolites critical for redox balance and TCA cycle activity, including, (i) *SLC25A45*, associated with NMN (NAD^+^ precursor) transport, (ii) *SLC25A11*, the oxoglutarate/malate carrier critical for TCA cycle flux and redox shuttling by NADH transportation from cytosol (Fig 1c,f).^8,9^ To determine whether the concurrent upregulation of *SLC25A45* and *SLC25A11* reflects a compensatory response to *SLC25A51* suppression independent of *CDKN2A* status, we stratified both *CDKN2A*-deleted and *CDKN2A*-intact cases by *SLC25A51* expression (top and bottom 25%) and performed correlation analyses. Interestingly, in the *CDKN2A*-deleted cohort, *SLC25A51* expression negatively correlated with both *SLC25A11* and *SLC25A45* (r ≈ -0.3), whereas no significant correlation was observed in *CDKN2A*-intact tumors (Fig 1d, e), indicating that this compensatory rewiring is genotype-specific, rather than a general mitochondrial stress response. Furthermore, *SLC25A20*, the carnitine-acylcarnitine translocase essential for fatty acid β-oxidation driven OXPHOS was also upregulated in *CDKN2A*-deleted tumors (Fig 1c).^10^ Unlike published findings in *SLC25A51*-null cells, we did not observe differential expression of the mitochondrial folate/FAD carrier *SLC25A32* (Fig 1c).^5^

**Fig 1.**
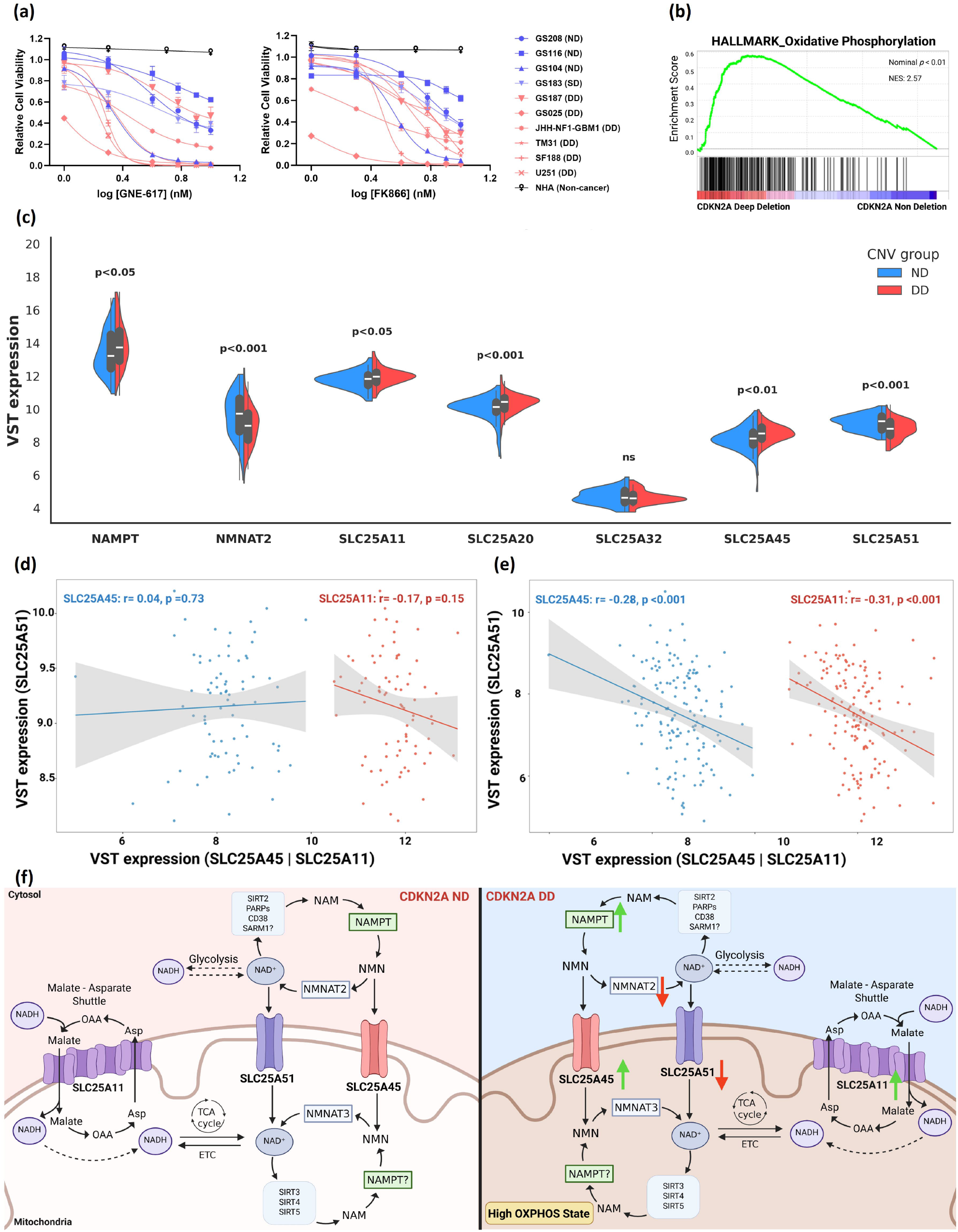
(a) Dose-response curves of GBM cell lines to NAMPT inhibitors GNE-617 and FK866. *CDKN2A*-deleted lines show heightened sensitivity than intact counterparts; NHA cells (non-cancer control) remain unaffected up to 100 nM. Data are mean ± SD (n = 4, ≥3 independent experiments, ND: Non-deletion, SD: Shallow-deletion, DD: Deep Deletion of *CDKN2A*); (b) GSEA of TCGA GBM data showing significant enrichment of mitochondrial oxidative phosphorylation signatures in *CDKN2A*-deleted tumors; (c) Variance stabilizing transformation (VST) expression of genes by *CDKN2A* status in TCGA-GBM: non-deletion (ND, blue; copy number ≥2) vs deep-deletion (DD, red; copy number = 0, homozygous deletion); (d-e) Gene-gene correlation of *SLC25A51* expression with *SLC25A45* and *SLC25A11* by *CDKN2A* status. Negative correlation observed in *CDKN2A*-deleted tumors (r ≈ -0.3) (e); no significant correlation in *CDKN2A*-intact tumors (d); (f) Summary schematic of pathway and gene expression differences in *CDKN2A* DD and ND tumors (Created in BioRender. https://BioRender.com/bf73dzj).

## Conclusions

Together, these data support a model in which *CDKN2A*-deleted GBM cells sustain high OXPHOS despite reduced cytosolic NAD^+^ synthesis (lower *NMNAT2* expression) and constrained mitochondrial NAD^+^ import (lower *SLC25A51* expression) by compensating through alternative NADH/NMN shuttling via *SLC25A11* and *SLC25A45* upregulation. Consequently, they become acutely dependent on cytosolic NMN/NAD^+^ regeneration via NAMPT, explaining their heightened sensitivity to NAMPT inhibition. While our analyses provide robust correlative insights, we do acknowledge tumor heterogeneity as a limitation of transcriptome-based inferences. These preliminary findings support the emerging concept that considering both metabolic state and genetic context can improve therapeutic targeting in GBM. However, fully achieving this potential will require embracing and continuing to explore the complexities of synthesis, transport and catabolism of the NAD^+^ metabolome.

## Funding

This research was funded by the UCLA SPORE in Brain Cancer (P50CA211015).

## Conflict of Interest

None

## Author Contributions

SD: Conceptualization, experiment conduction, data analysis, manuscript writing, and figure preparation. GY: Computational analyses. RAH: Data calculations and graphic preparation. CT: Technical assistance and experimental support. DAN: Provided resources and reviewed the manuscript. AL: Provided resources. KV: Provided resources. FJR: Manuscript review and project oversight. All authors contributed to the article and approved the submitted version.

## Data Availability

Data generated as part of this project will be made available upon request.

## Ethics Statement

All experimental procedures were approved by the Institutional Review Board committee of UCLA.

## Notes

https://github.com/EdgeYu97/CDKN2A-GBM-Analysis

### Competing Interest Statement

The authors have declared no competing interest.

## References

1. Shibao S, Minami N, Koike N, Fukui N, Yoshida K, Saya H, & Sampetrean O. Metabolic heterogeneity and plasticity of glioma stem cells in a mouse glioblastoma model. Neuro-oncology. 2018; 20(3): 343–354. DOI: 10.1093/neuonc/nox170

2. Gujar AD, L. S, Mao DD, Dadey DY, Turski A, Sasaki Y, et al. An NAD^+^-dependent transcriptional program governs self-renewal and radiation resistance in glioblastoma. Proceedings of the National Academy of Sciences. 2016; 113(51): E8247–E8256. DOI: 10.1073/pnas.1610921114

3. Redler J, Nelson AE, Heske CM. Mechanisms of resistance to NAMPT inhibitors in cancer. Cancer Drug Resistance. 2025; 8: 18. DOI: 10.20517/cdr.2024.216

4. Luongo TS, Eller JM, Lu MJ, Niere M, Raith F, Perry C, et al. SLC25A51 is a mammalian mitochondrial NAD^+^ transporter. Nature. 2020; 588(7836): 174–179. DOI: 10.1038/s41586-020-2741-7

5. Kory N, uit de Bos J, van der Rijt S, Jankovic N, Güra M, Arp N, et al. MCART1/SLC25A51 is required for mitochondrial NAD transport. Science advances. 2020; 6(43): eabe5310. DOI: 10.1126/sciadv.abe5310

6. Chen YJ, Iyer SV, Hsieh DCC, Li B, Elias HK, Wang T, et al. Gliocidin is a nicotinamide-mimetic prodrug that targets glioblastoma. Nature. 2024; 636(8042): 466–473. DOI: 10.1038/s41586-024-08224-z

7. Zacksenhaus E, Shrestha M, Liu JC, Vorobieva I, Chung PE, Ju Y, et al. Mitochondrial OXPHOS induced by RB1 deficiency in breast cancer: implications for anabolic metabolism, stemness, and metastasis. Trends in cancer. 2017; 3(11): 768–779. DOI: 10.1016/j.trecan.2017.09.002

8. Chen L, Wang P, Huang G, Cheng W, Liu K, Yu Q. Quantitative dynamics of intracellular NMN by genetically encoded biosensor. Biosensors and Bioelectronics. 2025; 267: 116842. DOI: 10.1016/j.bios.2024.116842

9. Lee JS, Lee H, Lee S, Kang JH, Lee SH, Kim SG, et al. Loss of SLC25A11 causes suppression of NSCLC and melanoma tumor formation. EBioMedicine. 2019; 40: 184–197. DOI: 10.1016/j.ebiom.2019.01.036

10. Yuan P, Mu J, Wang Z, Ma S, Da X, Song J, et al. Down-regulation of SLC25A20 promotes hepatocellular carcinoma growth and metastasis through suppression of fatty-acid oxidation. Cell Death & Disease. 2021; 12(4): 361. DOI: 10.1038/s41419-021-03648-1

